# Resumption of spermatogenesis in senescent goldfish *Carassius auratus* (Linnaeus, 1758) through spermatogonial cell therapy

**DOI:** 10.1101/428102

**Authors:** Sullip Kumar Majhi, Labrechai Mog Chowdhury, Santosh Kumar, Rajeev Kumar Singh, Vindhya Mohindra, Kuldeep Kumar Lal

**Affiliations:** ICAR-National Bureau of Fish Genetic Resources, Division of Fish Conservation. Canal Ring Road, Dilkhusa P.O., Lucknow 226 002, Uttar Pradesh, India

**Keywords:** Senescence, Reproduction, Spermatogenesis, Progeny, Goldfish

## Abstract

In recent times, stem cell research has gained considerable prominence because of its applications in assisted reproductive technology and the treatment of deadly diseases. In teleost fishes, spermatogonial stem cells have been effectively used to produce progeny of difficult-to-breed fish species and/or commercially valuable species through the surrogacy technique. The present study is the first report of an innovative application of stem cell therapy in teleostean fish species for revitalising the reproductive competence of senescent individuals. Senescent male goldfish, *Carassius auratus* aged approximately 10 years were procured from an ornamental fish-breeding farm and were reared locally for an additional 2 years. The senescence of the fish was evaluated and confirmed using histological analysis, gonadal index assessment, and germ-cell specific *vasa* gene expression. Analyses revealed the absence of spermatogonial cells and other germ cells in the testes of the senescent fish (n = 5). Spermatogonial cells from a prepubertal *C. auratus* male donor were isolated using discontinuous percoll gradients, labelled with the fluorescent dye PKH-26, and transplanted into the gonads of senescent *C. auratus* males through the urogenital papilla. Six months after the therapy, spermatozoa from males were collected through applying gentle manual pressure on the abdomen and were observed under the microscope. All the senescent therapy-treated *C. auratus* males produced spermatozoa from the transplanted cells; this was confirmed by retention of PKH-26 in the spermatozoa and diagnostic SSR locus. The senescent males were crossed with gravid *C. auratus* females through artificial insemination and natural spawning, and viable progeny was produced. These observations suggest that the reproductive competence of senescent individuals of commercially valuable and/or endangered fish species can be revitalised and extended through spermatogonia stem cell therapy to produce functional gametes.

## Introduction

Stem cells are described as undifferentiated cells that have the potential to renew themselves and differentiate into a single cell type or multiple specialised cell types [1]. This implies that they can undergo numerous cycles of cell division while maintaining their undifferentiated state and they have the capacity to differentiate into specialised cell types. Currently, stem cell research is one of the upcoming areas of research in science. Since their discovery and the subsequent establishment of protocols for successful isolation and culture [2] researchers have used them for various purposes, including treatment of chronic diseases and reproductive ailments [3]. Stem cell therapy has also shown the potential to revitalise or repair malfunctioning organs by using unscathed donor cells, which can take on the functions of damaged organs and provide animals with an extended and improved quality of life [4]. In humans, this has opened new avenue for regenerative medical approaches and has become a viable treatment option for healing chronically damaged organs [5]. However, in teleost fishes, the use of stem cells for therapeutic purposes has not been explored thus far; nevertheless, this therapy has the potential to revitalise malfunctioning reproductive organs of both sexually incompetent or senescent fishes that have a high commercial value or have become critically endangered. Currently, in fishes, stem cells are widely used for surrogate brood stock development through germ-cell transplantation [6–8]. In this case, donor cells obtained from young hatchlings or adults are transplanted into the recipient fish at various developmental stages, and on maturation, the recipient fish produce donor-derived gametes [8,9]. In this study, for the first time, we demonstrated successful restoration of spermatogenesis in senescent *C. auratus* through intragonadal stem cell therapy. The simple approach described in this study ultimately leads to the generation of viable functional spermatozoa from senescent *C. auratus* males, which can fertilise eggs derived from young *C. auratus* females and produce healthy progeny; thus, this technique extends the reproductive lifespan of fish beyond the pubertal phase and effectively generates viable progeny for aquaculture.

## Materials and Methods

### Ethics Statement

This study was approved by the Animal Ethics Committee of ICAR-National Bureau of Fish Genetic Resources. All the fish used in the experiments were handled according to the prescribed guidelines. During the study, the fish were sacrificed using an anaesthetic overdose and the gonads were excised.

### Animals, Age, and Rearing Protocols

Male goldfish (C. *auratus)* (n = 60; mean body weight ± standard deviation [SD] of 230 ± 20.5 g) were procured from Maharashtra, India and reported to be approximately 10 years old by the providers. The age of the fish was evaluated and confirmed by counting the growth rings located at scale (the growth rings from the farm raised fish with known age was used as reference). The fish were stocked in 500L tanks at a density of 5.0 kg of fish per m^3^ and reared in flow-through fresh water system (Temperature: 25°C ± 2°C; dissolved oxygen: 5.3–6.1 ppm; pH: 7.5–8; hardness: 40–45 ppm) under a constant light cycle (12 h light and 12 h dark). The fish were reared for an additional 2 years prior to the confirmation of their senility through histological analysis and germ-cell-specific *vasa* gene expression studies. The prepubertal donor *C. auratus* males (2–3 months old) were produced at the rearing facilities of the National Bureau of Fish Genetic Resources, Lucknow. Both groups of animals were fed a pelleted commercially available diet twice per day to satiation.

### Gonad Histology

After the completion of 2 years of rearing, five fish were randomly sampled and sacrificed using an overdose of anaesthesia (2-phenoxyethanol; Himedia, India), and their body weights were recorded. The gonads of the fish were excised and weighed, macroscopically examined, and photographed using a digital camera. The middle portion of the right and left gonads from each fish were then immersed in Bouin’s fixative for 24 h and preserved in 70% ethanol. The gonads were processed for microscopic examination by using routine histological procedures up to the stage of preparing 5-μm thick cross-sections and staining them with hematoxylin–eosin. Serial histological sections from each fish were examined under a microscope at magnifications between 10× and 60×.

### Gene Expression Analysis

Samples for real-time reverse transcription polymerase chain reaction (RT-PCR) of *vasa* gene expression were obtained from the anterior region of the testes after the completion of the 2-year rearing period and at 6 months after spermatogonial cell therapy; the samples were stored in RNA*later* (Sigma–Aldrich, St.Louis, MO, USA) at −80°C until further processing. RNA was extracted using TRIzol (Invitrogen Life Technology, Carlsbad, CA, USA) according to the manufacturer’s protocol. cDNA was synthesised using a first-strand cDNA synthesis kit (Thermoscientific, USA). The primers for real-time RT-PCR were (5’-AACCCTCATGTTCAGCGCCAC-3’ and 5’-TGGTTTCAACAAAGACCATCGTGC-3’). The real-time PCRs were run in an ABI PRISM 7300 (Applied Biosystems, USA) system using *Power* SYBR^®^ Green PCR Master Mix in a total volume of 15 μL, which included 7.5 μL of 2× Maxima™ SYBR Green qPCR Master Mix (Thermoscientific, USA), 20 ng of first-strand cDNA and 5 pmol L^−1^ of each primer. *β*-actin (5’-GAC TTC GAG CAG GAG ATG G-3’ and 5’-CAA GAA GGA TGG CTG GAA CA-3’) was used as an endogenous control. The comparative Ct method was used for *vasa* mRNA quantification. The fold change in the expression level was calculated using the 2–ΔΔCt method [10].

### Isolation and Labelling of Donor Cells

The prepubertal donor *C. auratus* males (n = 3) were sacrificed using an anaesthetic overdose, and the testes were excised and rinsed in phosphate-buffered saline (PBS; pH = 8.2). The testicular tissue was finely minced and incubated in a dissociating solution containing 0.5% trypsin (pH 8.2; Worthington Biochemical Corp., Lakewood, NJ), 5% fetal bovine serum (JRH Biosciences, Lenexa, KS), and 1mmol L^−1^ Ca^2+^ in PBS (pH 8.2) for 2 h at 22°C. The dispersed testicular cells were sieved through a nylon screen (mesh size 50 μm) to eliminate the non dissociated cell clumps; suspended in discontinuous percoll (Sigma–Aldrich, St. Louis, MO, USA) gradients of 50%, 25%, and 12%; and centrifuged at 200 ×*g* for 20 min at 20°C [8]. The phase containing predominantly spermatogonial cells (determined during preliminary trials which involved cell size measurement) was harvested, and the cells were subjected to rinsing as well as a cell viability test through the trypan blue (0.4% w/v) exclusion assay. The cells were then exposed to the PKH-26 Cell Linker (Sigma–Aldrich, St.Louis, MO, USA) at a concentration of 8 μmol/mL (room temperature, 10 min) to label the cells for tracking their behaviour inside the senescent (recipient) *C. auratus* gonads. The staining procedure was stopped by the addition of an equal volume of heat-inactivated fetal bovine serum. The labelled cells were rinsed three times to remove the unincorporated dye, suspended in Dulbecco Modified Eagle Medium (Life Technologies, Rockville, MD) with 10% fetal bovine serum, and placed on ice until transplantation.

### Cell Therapy Procedure

Forty fish were first anaesthetised using 200 ppm phenoxyethanol (Himedia, India) and placed on a cell transplantation platform, where they received a constant flow of oxygenated water containing 100 ppm of the anaesthetic drug. To prevent desiccation, the surface of the fish was moisturised during the entire procedure of cell therapy, which lasted for approximately 5–7 min per fish on average. A micro syringe was used to inject the cell suspension into the testicular lobe through the genital papilla (Fig. 1). Each fish was injected with 100 μL of a cell suspension containing approximately 5 × 10^4^ cells, at a flow rate of approximately 20 μL/min. Trypan blue was added to the injection medium prior to transplantation to facilitate visualisation of the cell suspension inside the needle and inside the gonads after injection. The genital papilla region was topically treated with 10% isodine solution and the fish were resuscitated in clean water.

**Figure 1.**
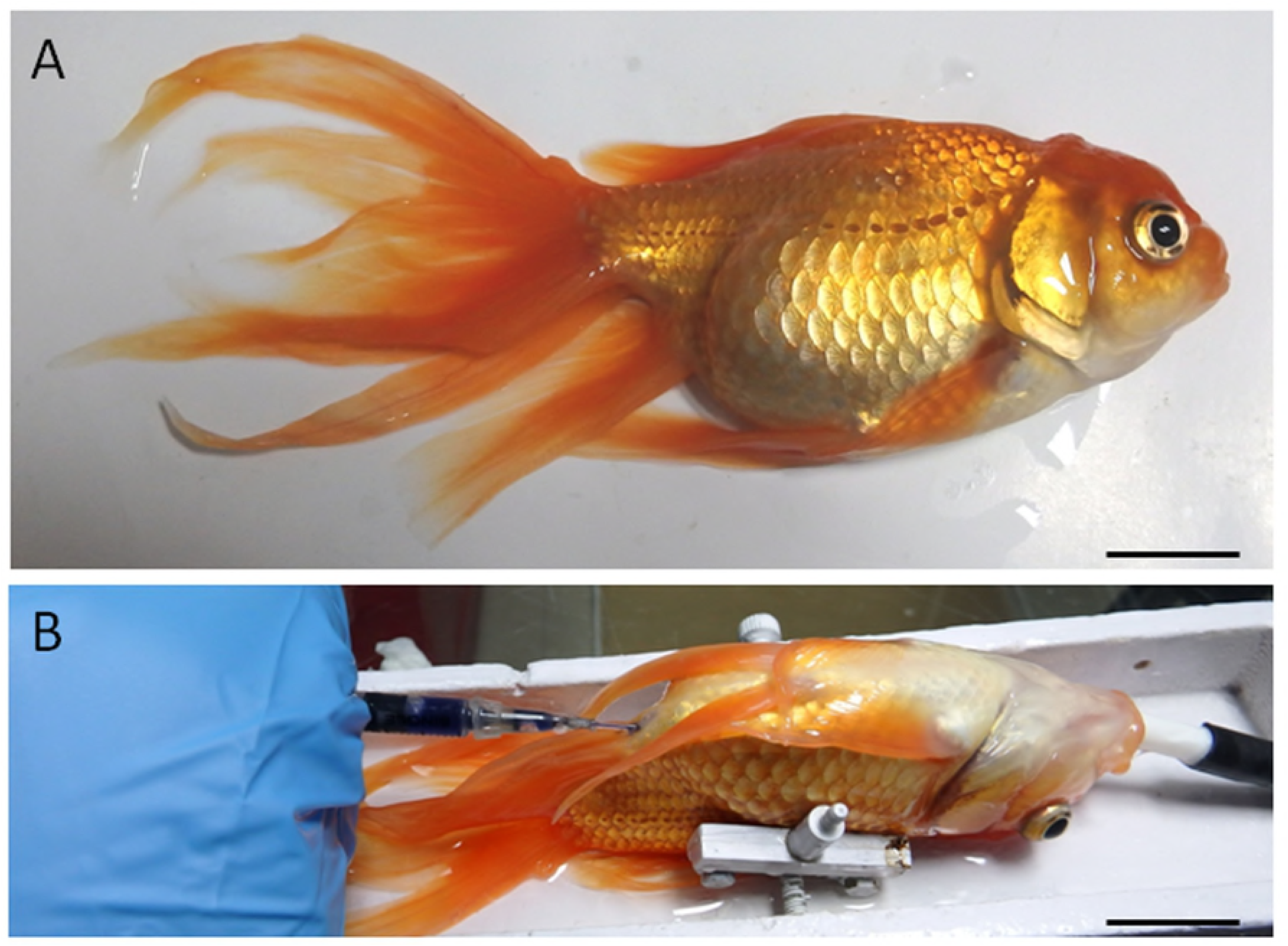
Model fish species used in this study (A: senescence *C. auratus* males) and the intra-papillar transplantation of donor cells into recipient gonads (B). The senescent recipients were placed on a platform and received a constant flux of aerated anaesthetic water through the gills during the procedure. The medium containing the donor cells was visualized by addition of Trypan blue during injection through the genital papilla. Scale bar indicates 2.5 cm.

### Analysis of donor cells Post therapy

The fate of donor cells after therapy was analysed through fluorescent microscopy at 6 and 12 weeks after injection. Hence, the testes from the five animals chosen randomly during each sampling were removed, washed in PBS (pH 8.2), macroscopically examined for the degree of dispersion of the cell suspension (Fig. 2), and immediately frozen in liquid nitrogen. Cryostat (Leica CM 1500, Germany) sections of 8-μm thickness were cut from representative portions of these testes and observed under a fluorescent microscope (Nikon Eclipse E600, Tokyo, Japan) for detecting the presence of PKH-26-labelled donor cells.

**Figure 2.**
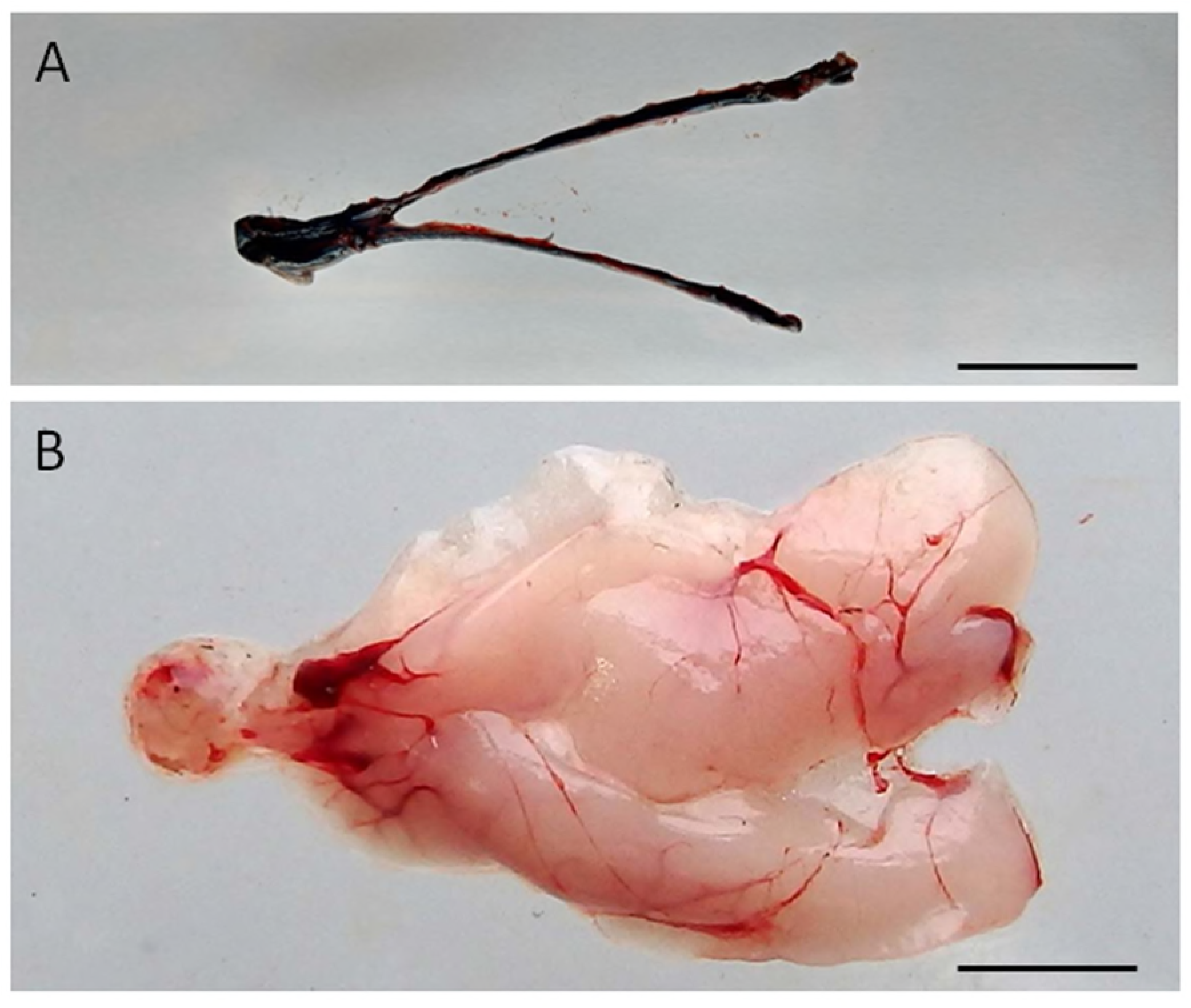
Visualization of the dispersal of the cell suspension through the gonad of senescence *C. auratus* after transplantation. A) Macroscopic appearance of the testis 48 hours after spermatogonia cell transplantation (note the diffusion of the marker Trypan blue throughout the testis). B) Appearance of the testis 6 months after cell transplantation (note the transplanted cells in senescence testis have undergone proliferation and differentiation to match the testis of a sexually mature individual). Scale bars indicate 2 cm.

The fate of the donor cells was then examined by measuring the gonado-somatic index (GSI; n = 5) and sperm density at 6 months after therapy. On each occasion, 10–30 μL of sperm was collected from each of the 25 therapy-treated males and 5 control males. Sperm was manually stripped by gently applying abdominal pressure after careful removal of urine and wiping the genital papilla. The sperm samples (10 μL) were then diluted 1,000 times with PBS and the density of spermatozoa was counted using a haemocytometer (Kayagaki Irika Kogyo Co., Ltd., Tokyo, Japan) under a microscope. Some of the spreads of sperm used for counting were also observed under the fluorescent microscope for the detection of PKH-26-labelled cells.

Sperm and blood samples were collected at 18 months from the spermatogonia cell transplanted senescent male. Total DNA was extracted using PureLink Genomic DNA kit (Invitrogen Life Technology, Carlsbad, CA, USA) according to the manufacturer’s protocol and DNA typing was done using *C. auratus* SSR locus *J60\** (forward 5’-CTGGCTGTCTGATCCTGCTGAT-3’ and reverse 5’-TGGCCAGAGTTTAAAAACCAGTCC -3’) [11]. PCR reaction was performed in 25μL consisted of 1x Taq buffer, 1.5 mM MgCl_2_, 0.5μm each primer, 200 μM of dNTP, 0.75 U Taq DNA polymerase and 80ng of DNA template. The amplification was done in an Applied Biosystem (Veriti) Thermal Cycler (ThermoFisher Scientific, USA) and consisted of an initial denaturation at 94°C for 5 min, 25 cycles of 94°C for 30 sec, 58°C for 30 sec and 72°C for 1 min, followed by elongation at 72°C for 4 min. Amplified products was visualized by 10% polyacrylamide gel electrophoresis (PAGE) and silver staining. Amplicon sizing was done on gel imaging and analysis system (UVP) using Msp I digested pBR322 as ladder.

### Artificial Insemination and Natural Spawning

After 6 months, the senescent therapy-treated *C. auratus* males resumed production of spermatozoa. Artificial insemination and natural spawning were performed using eggs from wild *C. auratus* females. Approximately 20 μL of milt from each of ten therapy treated males was used to fertilise a batch of *C. auratus* eggs; the batches of eggs were then incubated under flowing water at 25°C until hatching. In the natural spawning trials (see Supplementary Video), ten senescent therapy-treated males and wild females were paired in a 100L glass aquarium and reared in fresh water (Temperature: 25°C ± 2°C; dissolved oxygen: 5.3–6.1 ppm; pH: 7.5–8; hardness: 40–45ppm) under a constant light cycle (10 h light and 14 h darkness). Every, morning between 10:00 and 11:00 h, the tanks were examined for spawning and embryos were collected. The fertilised eggs obtained from each cross were incubated at 25°C and observed under a light microscope for the assessment of fertilisation, embryonic development, and hatching.

### Statistical Analysis

The statistical significance of the differences in *vasa* gene expression levels, GSI values, and sperm densities among the groups was evaluated using one-way analysis of variance and Tukey’s multiple comparison test. Graphpad Prism ver. 4.00 (Graphpad Software, San Diego, Carlifornia, USA) was used for statistical analyses. Data are presented as mean ± SD, and the differences among the groups were considered as statistically significant at *P* < 0.05.

## Results

### Histological Observation of Germ Cells in Senescent Males

Microscopic examination of the testes revealed that all the five senescent *C. auratus* males aged 12 years exhibited shrunken gonads (Fig. 2A) and complete disappearance of all the stages of germ cells (Fig. 3A-B) in all sections examined. By contrast, the control males (age: >1 year) exhibited active spermatogenesis with large cyst of spermatogonial cells, and all the stages of germ cells (Fig. 3C-D). The efferent duct was found to contain spermatozoa. These histological observations were also corroborated by the results of GSI evaluation and real-time RT-PCR analysis, which showed that both GSI (Fig. 4) values and *vasa* gene transcript levels (Fig. 5) were significantly lower in the senescent males than in the controls (P < 0.05).

**Figure 3.**
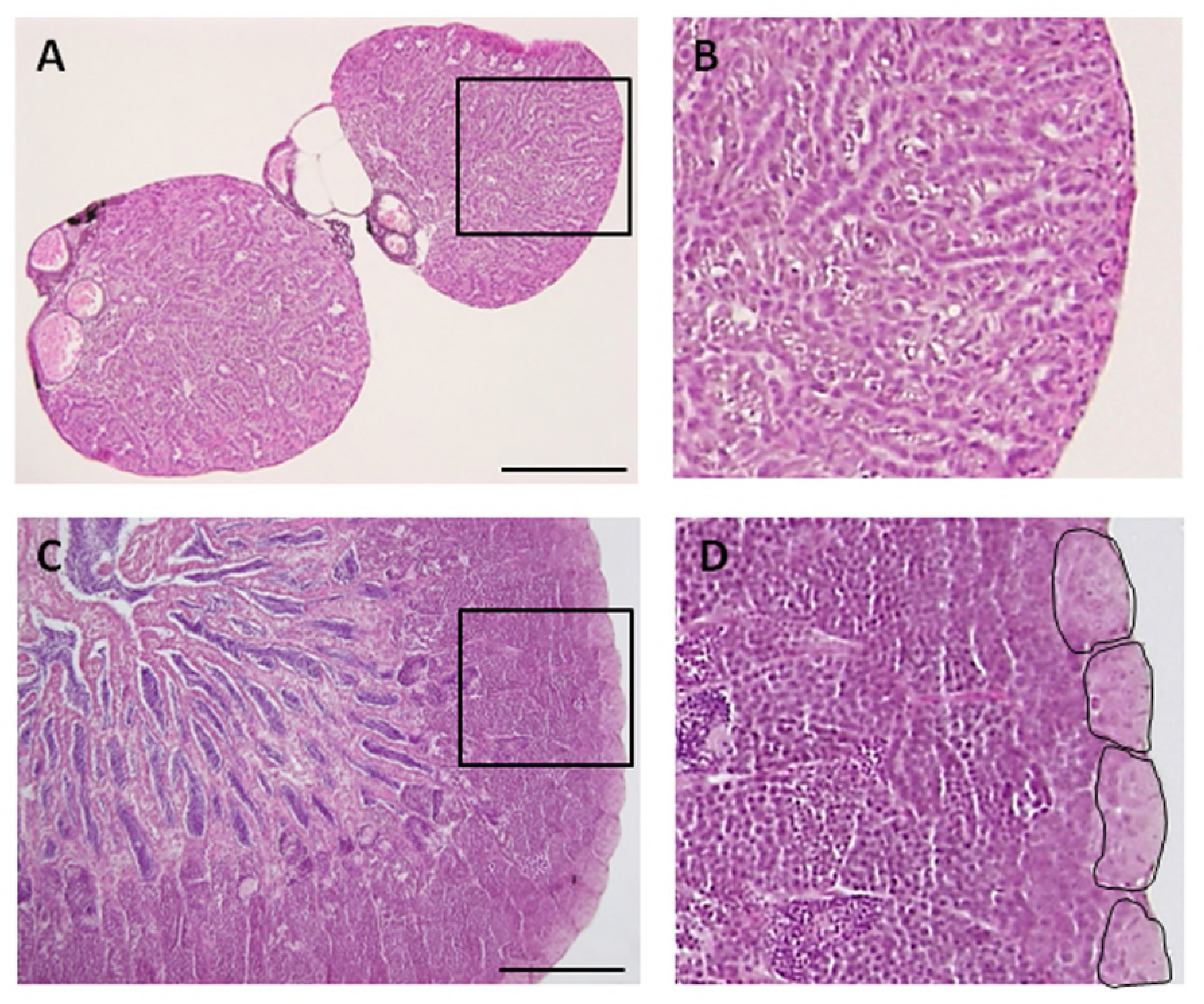
Histological appearance of senescence and control *C. auratus* gonads. Panels on the right are high magnifications of insets in the left panels. A,B) Twelve year old senescence testis showing complete absence of spermatogonia and other germ cells. C,D) A sexually mature control testis indicating large cysts of spermatogonia in the blind end of the spermatogenic lobules (highlighted) and an active spermatogenesis within the lobules. Scale bars indicate 100 μm (A and C) and 20 μm (B and D).

**Figure 4.**
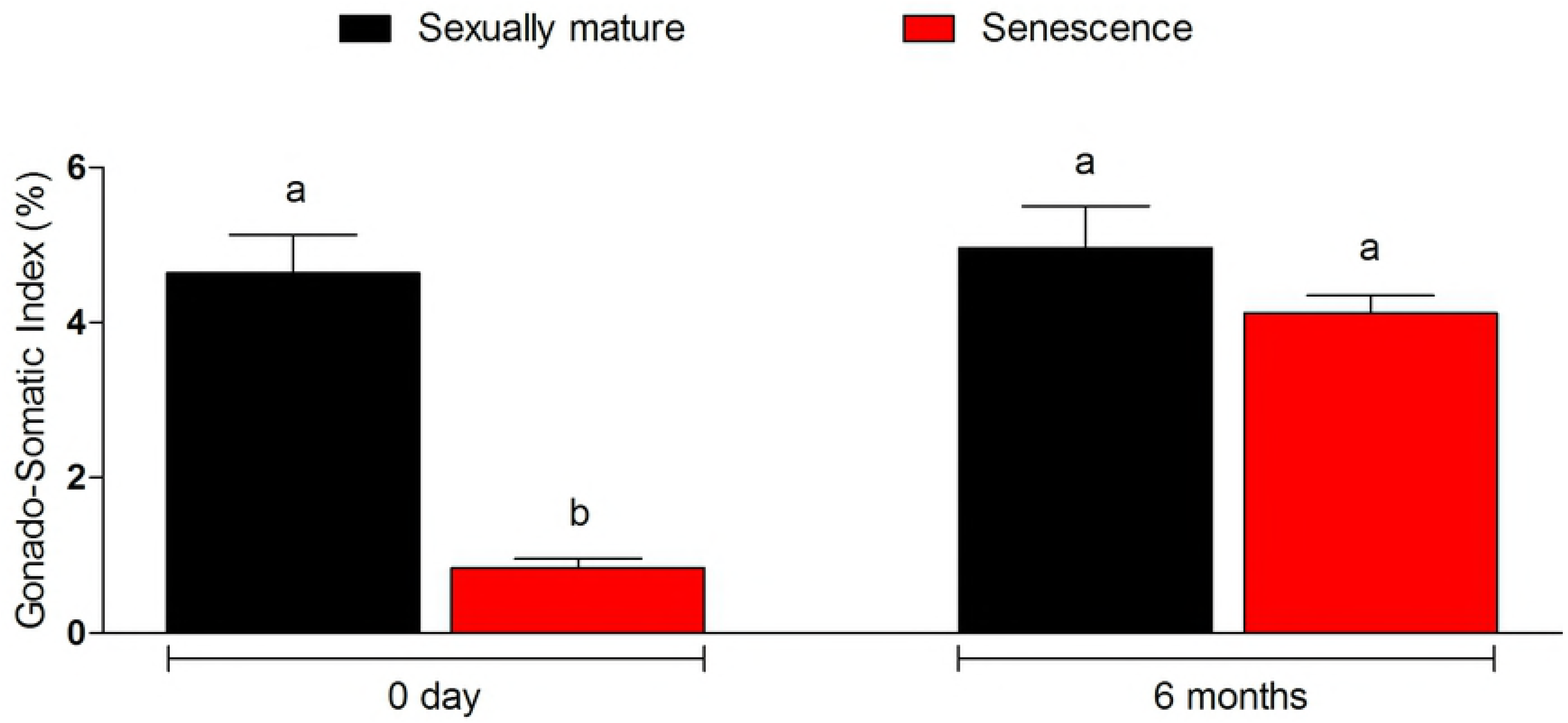
Changes in the gonado-somatic index of senescence and control gonads between 0 and 6 months. Note, 6 months after therapy the gonado-somatic index value of senescence *C. auratus* males have significantly increased to match the sexually mature control. Columns with different letters vary significantly (ANOVA - Tukey test, *P*<0.05).

**Figure 5.**
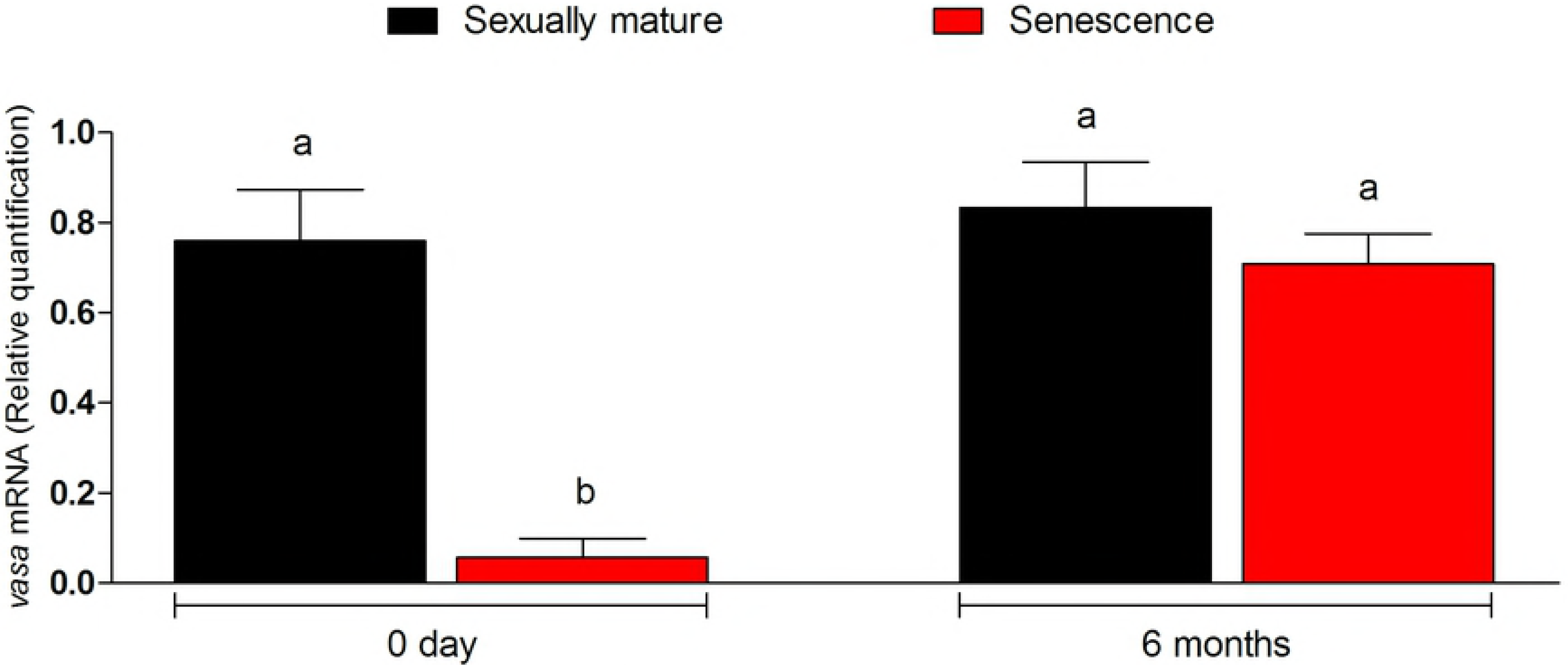
Changes in *vasa* gene transcript levels in senescence and control C. *auratus* gonads between 0 and 6 months. Note, 6 months after therapy the *vasa* gene transcript levels in senescence C. *auratus* males have significantly increased to match the sexually mature control. Columns with different letters vary significantly (ANOVA - Tukey test, P>0.05).

### Fate of Transplanted Cells in Senescent Testes

The transplanted donor spermatogonial cells were found randomly distributed throughout the spermatogenic lobules in all the five senescent males examined 6 weeks after the therapy. At 12 weeks after the therapy, the donor spermatogonial stem cells had reached the blind end of the lobules (cortical region of the testis; Fig. 6A-D) and had undergone proliferation to form a network along the testicular lobule; this stage was observed in all the sampled fish (n=5; Fig. 6E). At 6 months after the therapy, the sperm could be collected by applying gentle pressure on abdomen in all the 25 senescent *C. auratus* males (Fig. 6F). The sperm density, which was not detectable before the therapy in the senescent *C. auratus* males, significantly increased after the therapy and was comparable with that of sexually mature control males (Fig. 7).

**Figure 6.**
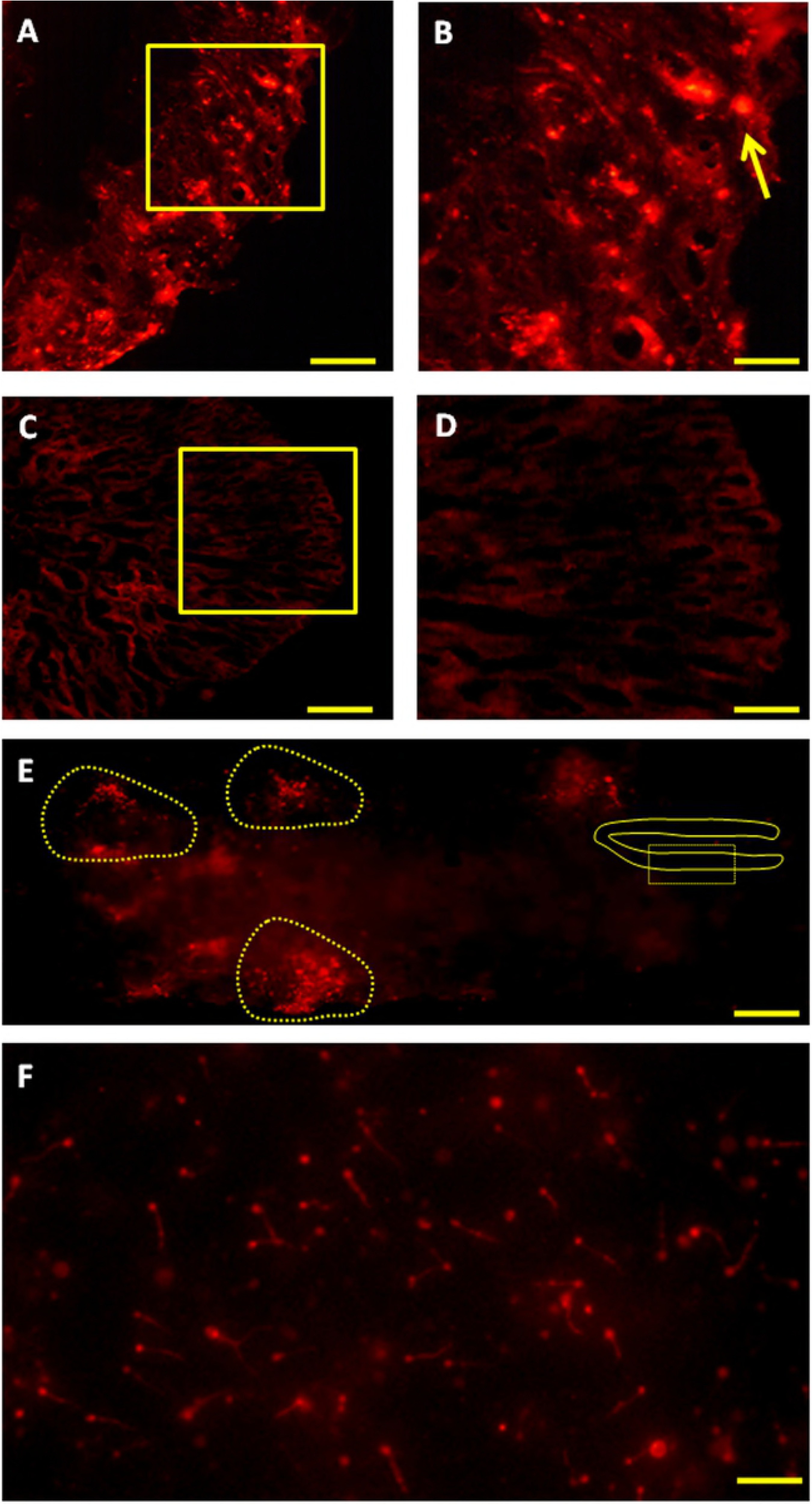
Fate of PKH-26-labeled donor spermatogonia cells in senescent *C. auratus* males examined between 4 weeks and 6 months after transplantation. A,B) Cryostat section of a transplanted testis at 4 weeks showing the presence of transplanted spermatogonia cell at the blind end of the spermatogenic lobules (arrow; B is a high magnification of the box in A). C,D) Cryostat section of a non-transplanted, control testis at 4 weeks showing the approximate location of the blind end of the spermatogenic lobules (D is a high magnification of the box in C). E) Whole-mount preparation of a transplanted testis at 12 weeks showing the proliferation of donor-derived cells (highlighted) along the length of the gonad. F) Whole-mount preparation of spermatozoa cell derived from the senescent male 6 months after the procedure (Note, the spermatozoa produced were of donor-origin that was characterized by retention of PKH-26 dye). Scale bars indicate 100 μm (A and C), 20 μm (B and D) and 500 μm (E and F).

**Figure 7.**
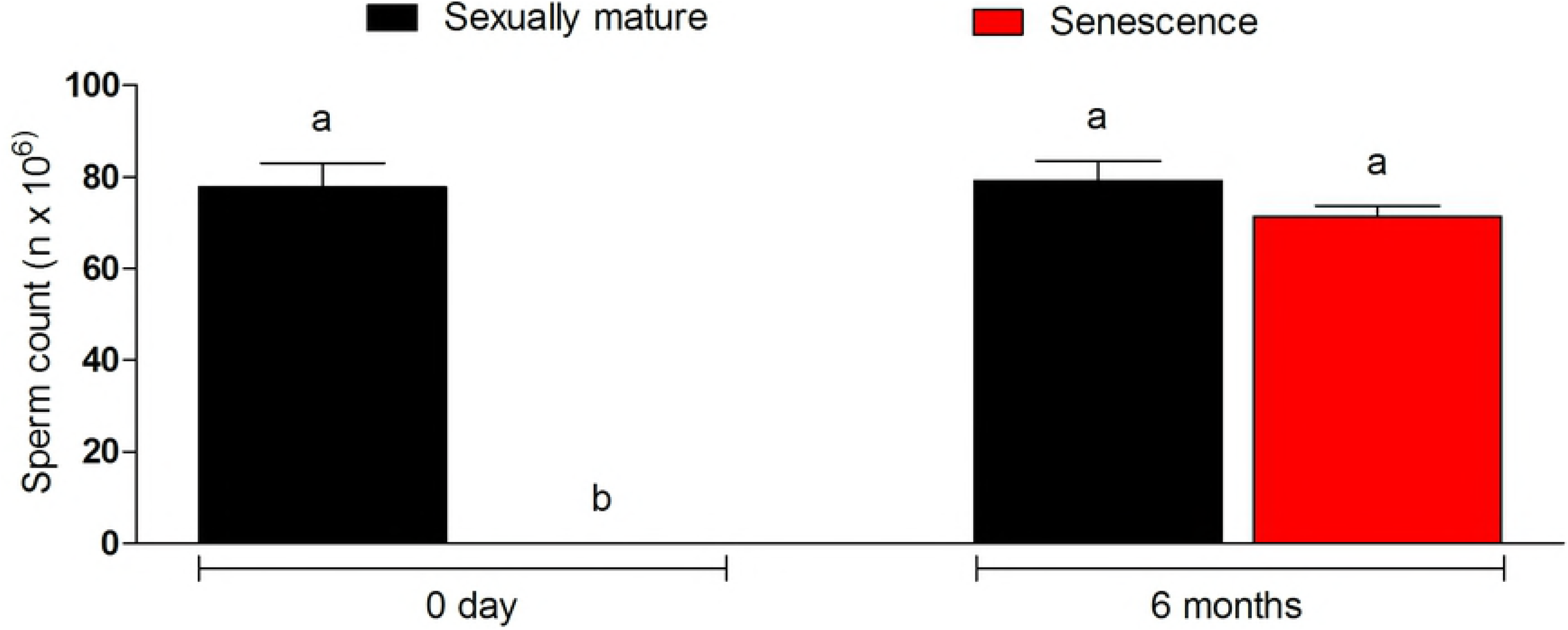
Sperm density in senescent recipients and non-transplanted (negative control) *C. auratus* between 0 and 6 months after spermatogonia cell transplantation. Note, the sperm count in the senescent males had significantly increased after spermatogonia cell therapy. Columns with different letters vary significantly (Tukey’s multiple comparison test, *P*<0.05).

### Production of Viable Gametes and Progeny from Senescent Males

Six months after cell therapy, the senescent *C. auratus* males were found to resume spermatogenesis and produce spermatozoa from the transplanted cells; the origin of the spermatozoa (from the transplanted cells) was confirmed by the existence of two different genotypes and presence of red fluorescent labels in the cells (Fig. 6F). Locus *J60\** differentiated amplicons from blood (recipient) and sperm (donor cell) DNA (Fig. 8). These therapy-treated males were then used for artificial insemination and for trials of natural spawning of eggs obtained from young *C. auratus* females (Tables 1–2). These crosses resulted in normal embryonic development and hatching, which were similar to those in the control animals. The crosses between the spermatozoa from therapy-treated males and eggs from wild *C. auratus* females resulted in 88.6%–97.5% hatching; oppose to 93.8%–97.7% in control (Table 1). In addition, when the senescent males were coupled with wild *C. auratus* females for natural spawning, the crosses resulted in 95.5%-99.5% hatching with normal embryonic development (Fig. 9) and viable progeny; oppose to 97.7%-98.4% in control (Table 2). These observations suggest the viability of the proposed approach in revitalising the reproductive competence of commercially valuable fish species that become senescent with age.

**Figure 8.**
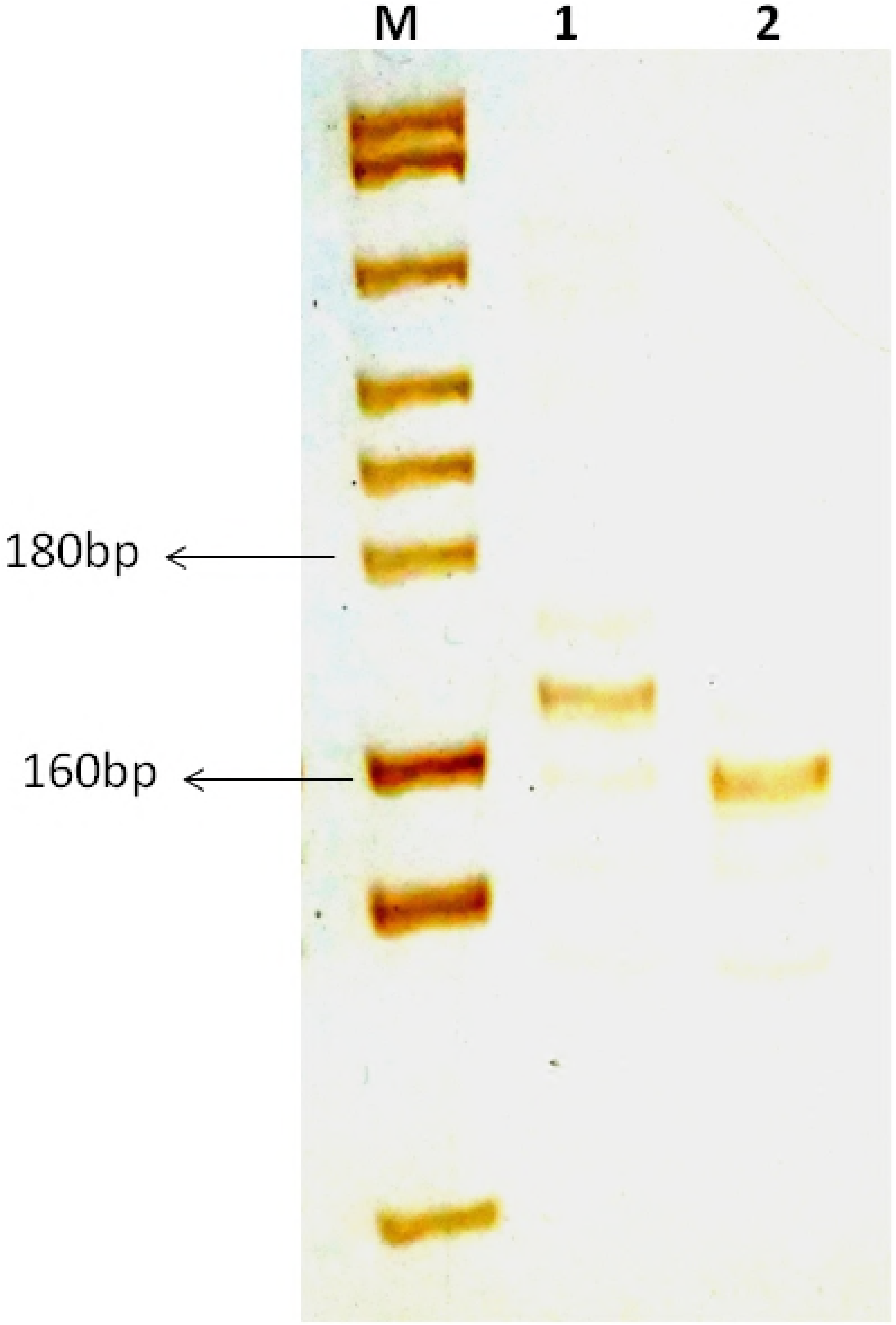
Genotyping of sperm and blood collected from a senescent *C. auratus* males 18 months after the cell transplantation. The microsatellite primers were used for amplification of DNA for genotyping. Lanes include molecular marker (M), blood of senescent male (1) and sperm (2). Note, the donor-derived spermatozoa were detected in the sperm of senescent recipients shown in lane 2.

**Figure 9.**
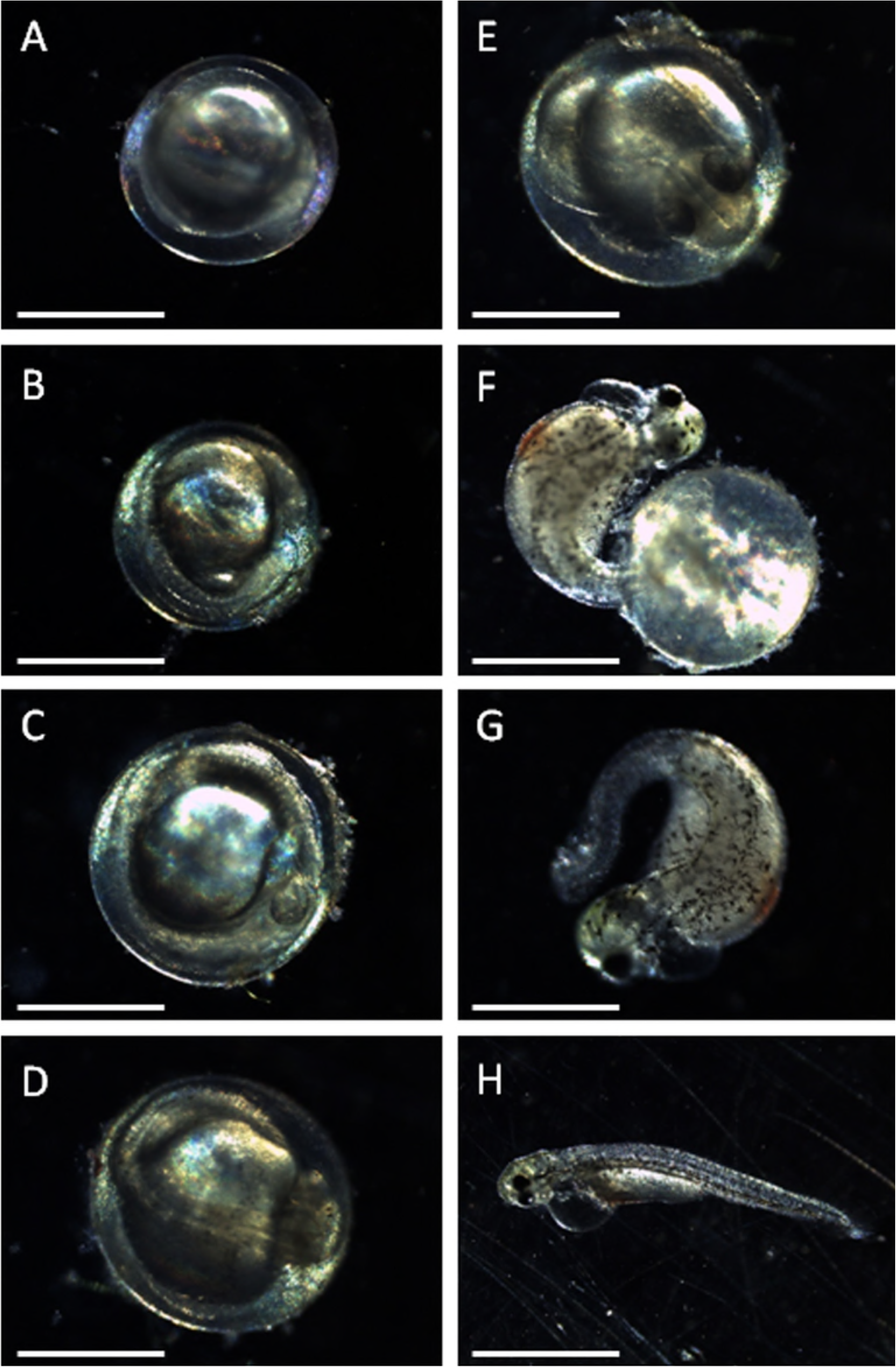
Embryos recovered from natural spawning (spermatogonia cell transplanted senescent *C. auratus* male paired with young wild female). Note that the embryos were reared at 25°C water temperature for hatching. A) Germ ring stage of embryo recovered at 7 hours 30 minutes after fertilization. B) At 12 hours after the fertilization, the embryo attained the bud stage. C) 6-somite stage was observed at 14 hours, 16-somite at 18 hours (D), and 22-somite stage at 22 hours after the fertilization (E). F) The pharyngula stage was observed at 44 hours. G) Long pac stage was observed at 58 hours and all the larvae hatched out 72 hours after fertilization (H) and exhibited active vertical and horizontal movement similar to progeny derived from control parents. Scale bar-500 μm (A-H).

**Table 1.** Results of artificial insemination of eggs derived from wild gravid *C. auratus* females with sperm from ten senescence *C. auratus* males (#1~10; transplanted with spermatogonia derived from pre-pubertal *C. auratus* donors) and control (crosses between sexually mature *C. auratus* males and females).

| senescence *C. auratus*<br>males | Eggs (n) from wild gravid<br>*C. auratus* females | Fertilization rate<br>(n; %) | Hatching rate<br>(n; %) |
| --- | --- | --- | --- |
| #1 | 430 | 411 (95.5) | 400 (97.3) |
| #2 | 610 | 586 (96.0) | 550 (93.8) |
| #3 | 360 | 327 (90.8) | 290 (88.6) |
| #4 | 550 | 530 (96.3) | 500 (94.3) |
| #5 | 470 | 455 (96.8) | 436 (95.8) |
| #6 | 510 | 492 (96.4) | 477 (96.9) |
| #7 | 630 | 610 (96.8) | 595 (97.5) |
| #8 | 280 | 243 (86.7) | 233 (95.8) |
| #9 | 440 | 415 (94.3) | 405 (97.5) |
| #10 | 485 | 465 (95.8) | 440 (94.6) |
| Control#1 | 615 | 580 (94.3) | 565 (97.4) |
| Control#2 | 425 | 405 (95.2) | 380 (93.8) |
| Control#3 | 470 | 445 (94.6) | 435 (97.7) |

**Table 2.** Results of natural spawning of cross between senescence *C. auratus* males (#1~10; transplanted with spermatogonia derived from pre-pubertal *C. auratus* donors) with wild gravid *C. auratus* females and a control (paired between sexually mature *C. auratus* males and females).

| senescence *C. auratus* males paired with gravid *C. auratus* females | Total number of embryo collected (n) | Fertilization rate (n; %) | Hatching rate (n; %) |
| --- | --- | --- | --- |
| #1 | 980 | 970 (98.9) | 945 (97.4) |
| #2 | 1020 | 996 (97.6) | 960 (96.3) |
| #3 | 1110 | 1087 (97.9) | 1072 (98.6) |
| #4 | 895 | 881 (98.4) | 865 (98.1) |
| #5 | 1500 | 1485 (99.0) | 1478 (99.5) |
| #6 | 1460 | 1448 (99.1) | 1438 (99.3) |
| #7 | 630 | 618 (98.0) | 598 (96.7) |
| #8 | 1220 | 1205 (98.7) | 1185 (98.3) |
| #9 | 810 | 785 (96.9) | 773 (98.4) |
| #10 | 1060 | 990 (93.3) | 946 (95.5) |
| Control#1 | 1245 | 1205 (96.7) | 1186 (98.4) |
| Control#2 | 970 | 951 (98.0) | 934 (98.2) |
| Control#3 | 1060 | 1045 (98.5) | 1022 (97.7) |

## Discussion

In the present study, donor spermatogonial cells harvested from prepubertal *C. auratus* testes were successfully used for restoring the reproductive competence of senescent *C. auratus* males by using cell-therapy intervention. As observed in mammals [12,13], the therapy resulted in recolonization of the seminiferous epithelium, resumption of spermatogenesis, and production of functional spermatozoa. The simple but effective procedures demonstrated in this study can be immediately applied in hatcheries and other seed production facilities. This procedure has considerable potential for extending the reproductive phase of commercially valuable species.

In the present study, the testes recovered from the senescent *C. auratus* males exhibited the shrinkage of the testicular lobes, absence of germ cells, and deposition of adipose and connective tissue. The germinal epithelium was continuous, and the testicular cross-section was noticeably smaller than that observed in sexually mature animals. Notably, this observation was contrary to that in senescent guppy (*Lepistes reticulatus*) in which the histological changes in the testes revealed a progressive increase in the percentage of lobules containing spermatids and spermatozoa and an increase in the deposition of melanin as well as adipose and connective tissue [14]. However, in senescent mosquito fish (*Gambusia affinis*) and European bitterling (*Rhodeus amarus*), partial testicular degeneration was observed to occur with a considerable reduction in the number of germ cells [15–17]. According to Finch, 1990 [18] such variations in the gonadal germ-cell profiles and testicular morphologies of senescent fishes are due to their growth patterns and origins of habitation. Thus, the characteristics of senescence in fishes cannot be generalised and vary widely depending on their habitation history and other physico-chemical parameters of their ecosystem [19]. For instance, tropical fish species grow faster and attain senescence comparatively earlier than do temperate fish species [20]. The experimental fish used in this study were reared in a tropical environment (temperature range: 25°C–32°C) for 10 years; probably this may explain why germ-cell degeneration in the senescent individuals was considerably more severe than that previously reported in other fish species.

Regardless of the differences between sterile (natural or experimentally induced) and senescent testes (in this case), the process of recolonization by the transplanted spermatogonial cells was similar to the previously reported process [6–8,21]. For instance, after cell therapy, many PKH-26-labelled spermatogonial cells eventually settled along the blind end of the seminiferous lobules and began proliferation within weeks of transplantation. Considering that only spermatogonial stem cells can migrate and settle at the basement membrane and resume the process of spermatogenesis [22], we can surmise that the cells used for therapy were spermatogonial stem cells. This conclusion is also proved by the fact that when the senescent *C. auratus* males (26 months after cell therapy) were repeatedly paired with wild *C. auratus* females for natural spawning, they continued to produce viable progeny to date. Transmembrane protein molecules present at the junctional complex located in the seminiferous tubules have been reported to transduce signals, maintain cell polarity, and mediate germ-cell migration [23]. Although we did not examine the endocrine regulation involved in the migration, proliferation, and differentiation of the donor spermatogonial cells inside the gonads of senescent *C. auratus* males, the results obtained in the present study (time-course observations of donor spermatogonial cells after therapy) indicated that the donor spermatogonial cells might have sensed and responded to the molecules released from the blind end of the seminiferous lobules in therapy treated senescent *C. auratus* males. Consequently, these spermatogonial cells underwent migration and settled down near the basement membrane and resumed spermatogenesis. However, the duration for which the *C. auratus* males treated with spermatogonial cell therapy retain their reproductive competence and produce functional gametes remains to be determined.

The present study demonstrated that spermatozoa derived from senescent *C. auratus* males exhibited functional properties similar to those of the control spermatozoa in terms of fertilisation, embryonic development, hatching and no evidence of defective spermatozoa was observed. Defective spermatogeneis has been reported to occur most prominently when the donor and recipient animals are phylogenetically distant [24]. In this study, the spermatogonial cells derived from prepubertal *C. auratus* males were transplanted into senescent *C. auratus* males; thus the transplanted cells were probably immunologically compatible with the gonadal environment of the senescent *C. auratus* males. Moreover, the progeny produced using the spermatozoa derived from therapy-treated *C. auratus* males exhibited a similar growth pattern to that of the progeny produced from the control animals (data not shown). These observations suggest that the reproductive competence of senescent *C. auratus* males could be successfully revitalised using spermatogonial cell therapy. Currently, we are examining the feasibility of the cell-therapy approach in revitalising the reproductive competence of female fish.

In conclusion, the most remarkable achievement of this study is the production of functional spermatozoa from senescent *C. auratus* males, after spermatogonial cells derived from the prepubertal *C. auratus* donor were transplanted into the testes of the senescent males through the genital papilla. This approach, which to the best of the author’s knowledge has been validated for the first time in fish in this study, is a method to generate functional gametes for additional years after attainment of senility. This is a crucial development in the breeding of rare and/or commercially valuable fish species that are developed as brooders and require sizeable investments in terms of feed and health care. Such fish species are used for only a few years by the hatcheries for seed production and are invariably discarded when they lose their reproductive competence because of age.

## Acknowledgements

The authors express sincere thanks to the Director, ICAR-NBFGR, Lucknow for providing all necessary help for perform this work. The help provided by the field staff of the National Bureau of Fish Genetic Resources, Lucknow in rearing the fishes and collecting the samples is also sincerely acknowledged.

Supplementary Video. Natural spawning trail in a 100L glass tank involving spermatogonia cell transplanted *C. auratus* male and young wild females.

**Author Contributions:**
S.K.M. Conceived, designed and supervised the study, interpreted results and findings, and wrote the manuscript; L.M.C. Contributed to the genotyping of gametes produced from senescent goldfish; S.K. Contributed to spermatogonial cell isolation, transplantation and post transplantation analysis of cells; R.K.S Contributed to DNA typing of the gametes, interpreted results and findings; V.M. Interpreted results and findings, contributed to manuscript writing and revising; K.K.L. Contributed to the interpretation of results, read and edited the manuscript.

